# Emergent Collective Bats Oscillation Dynamics on Tree Branches

**DOI:** 10.64898/2026.05.23.727420

**Authors:** Sunny Kumar, Ousmane Kodio

## Abstract

The flying fox bats roost in large colonies, suspended upside-down with minimal grip efforts from tree branches that are exposed to environmental disturbances. In this study, we investigate the oscillation dynamics of bats hanging from tree branches under natural conditions with wind. Bats modulate their grips to control the oscillation during wind disturbances and actively transform their postures. Using field observations, we analyze the angular deformation, speed, and phase of individual and collective bats’ swaying motions in response to environmental perturbations. We observed the mechanical coupling-based synchronization of collective bat oscillations on a tree branch. To rationalize this new phenomenon of bats synchronization behavior, we perform a table-top experiment of a physical model using active oscillators and passive systems. This work could inform the design of bio-inspired suspension systems and contribute to our understanding of animal balance and collective behavior in unsteady and complex environments.

---

Classical collective behaviors in animals such as bird flocking [1], synchronous flashing of fireflies [2], fish schooling [3], and insect swarming [4, 5] are characterized by coordinated movement driven by direct sensory interactions such as vision, touch, acoustic, or chemical signaling [6]. These systems serve functions such as navigation, foraging, or predator evasion and rely on dynamic alignment and rapid response among individuals [7]. Bats roost on tree branches in large colonies, exhibiting group behavior [8, 9]. Many inanimate systems also exhibits collective synchronized behaviors [10–13].

Bats display a unique behavior of hanging upside down from elevated surfaces such as tree branches, caves, and human-made structures [14]. This inverted posture is enabled by a tendon-locking mechanism in their feet, allowing them to maintain a passive grip without continuous muscle exertion or fatigue [15–17]. The hind limbs of a bat adapt to cling rather than walking, making this posture energy-efficient and structurally stable, which offers an advantage for escape, as bats can drop directly into flight using gravity to gain initial momentum. The natural tilt of the head of a bat while it is hanging reduces circulatory and neurological stress by regulating blood flow [18]. The roosting at height ecologically provides safety from ground predators and allows early detection of approaching threats [19], which represents an evolutionary strategy combining mechanical efficiency, physiological safety, and ecological advantage [20–22] across diverse environments.

Here we study *Pteropus giganteus*, the flying fox bats, that are distributed throughout India, including both urban and rural areas. We observe the collective bats’ oscillation while roosting on tree branches, hanging upside down. Their swaying motion emerges from mechanical coupling through the branch, influenced by wind and body movements, creating a dynamic collective behavior. The static configurations are well studied [18], but the dynamic responses, the collective sway of bats during roosting, remain less understood, highlighting gaps in knowledge about stability and environmental adaptation. We investigate the physical model of bat oscillations using active oscillators and a passive 3D-printed bat system. Here, we report a new phenomenon of synchronization of bats oscillations through the mechanical coupling of tree branches.

## 1. Bats Roosting and Oscillation

Bats exhibit perching and collective roosting behavior on trees, and an individual bat (Fig. 1A, Movie 1) hangs upside down from a tree branch using its hind limbs. A group of bats’ distributed spatial arrangement suggests coordinated roosting hanging from the banyan tree canopy (Fig. 1B), and the triangulated network overlays illustrate the varying canopy areas for perching, with small and medium zones. The tree canopy architectures are globular, spreading, vase, and columnar, which shape bat roost organization, and heat maps reveal layered roosting structures and relative height distributions (fig. S1A-D). The bats distributed roost in trees in groups ranging from small to large numbers (N = 21 to 216 individuals), as illustrated by the box-and-whisker distribution plot (Fig. 1C) (details in materials and methods).

**FIG. 1.**
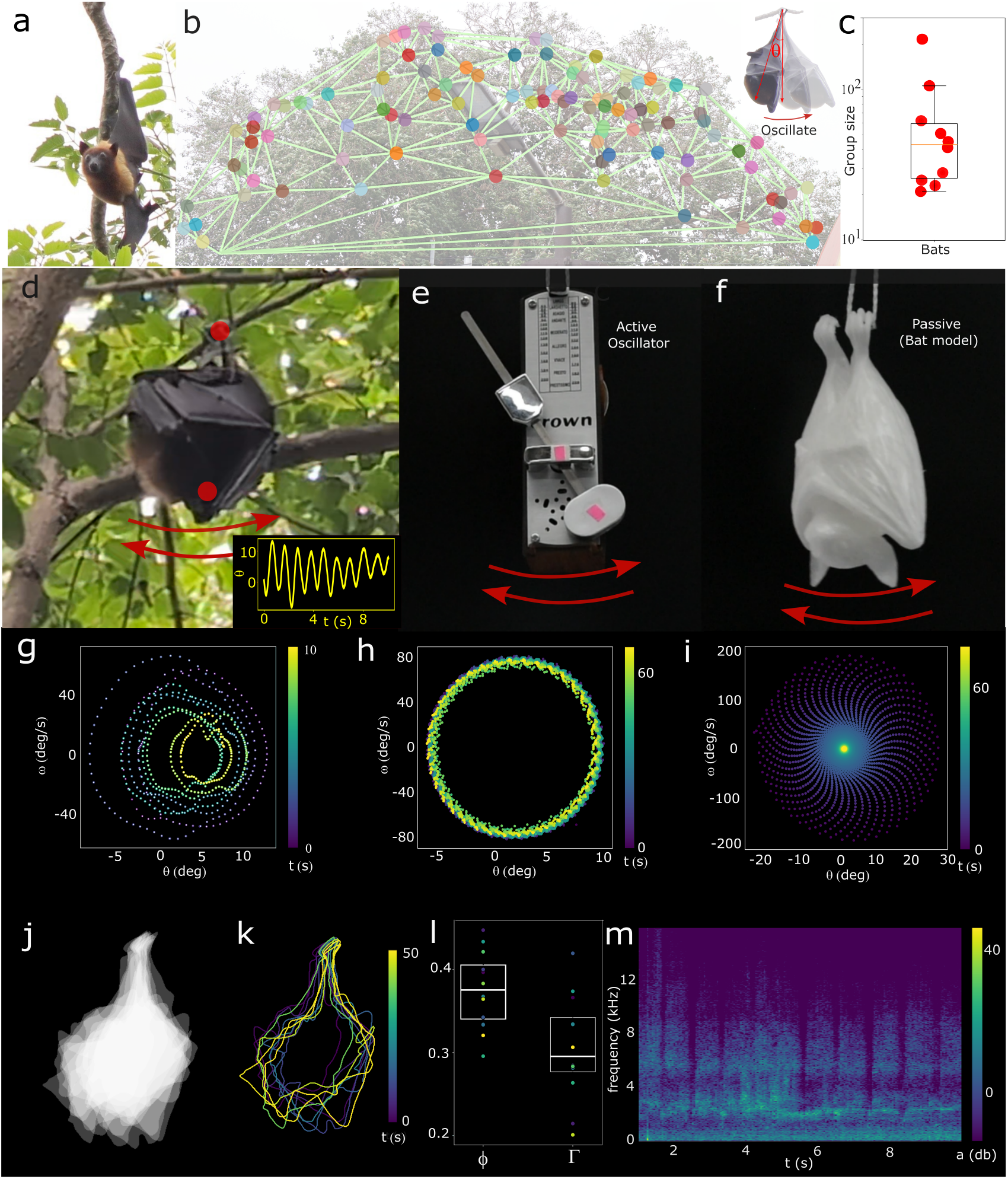
Spatial organization and oscillatory dynamics of bats on a tree branch. (**a**) An individual flying fox bat hanging upside down from a branch, with its hind limbs, depicts the passive-locking posture used during roosting. (**b**) Collective bats hanging and triangulated network overlays showing varying canopy areas used for perching zones, and the insert image shows angle, 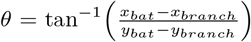 . (**c**) The box plots display the interquartile range, with the orange line indicating the median. The whiskers represent the data range, and the red dots denote individual bat group sizes on the tree. (**d**) Field photograph of a bat colony member roosting and swaying on a branch. The red dot marks indicate the tracking swaying point. (**e**) Physical active oscillatory model. (**f**) Physical passive 3D printed bat model. (**g**) Bat oscillation represents between phase space between theta, angular velocity 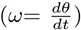. (**h**) A phase space between *θ, ω* for an active oscillation system. (**i**) A phase space plot for a passive 3D bat model. (**j**) Overlaid silhouettes of bats roosting on a branch, showing body posture distribution with time. (**k**) The bat shape transformation over time (color bar indicates progression from 0 to 50 s). (**l**) The box plots represent shape deformation of bats’ circularity (*ϕ* = 4*πA/P* ^2^) and deformability (Γ = (*a − b*)*/*(*a* + *b*)) between 50 s time intervals during hanging on a branch. (**m**) Spectrogram and waveform of bats’ roosting calls represented with frequency (*f*) with time and amplitude (a).

Individual bats swaying on a branch, where the trajectories of the swaying dynamics demonstrate the non-linear, multi-directional oscillations induced by external perturbation (Fig. 1D, fig S1, Movie 1). The bat oscillation period is *∼* 1–2 s, with a gradual decay in amplitude (energy loss) for the 10 s interval (fig. S1I). The angular deformation of a bat oscillation is represented by 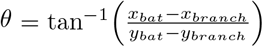, where (x_*bat*_,y_*bat*_) are the coordinates of the bat head position and (x_*branch*_,y_*branch*_) are the coordinates of the branch grip point. The bat oscillates with peak angular velocity showing 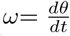 up to 60 deg/s and maximum acceleration 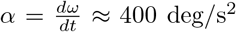 (Fig. 1 G, fig S1). The bat phase space (*θ, ω*) plot displays inward contraction of loops, confirming dissipative behavior, with the system progressively approaching equilibrium near the origin. This motion can be modelled as a simple pendulum system, where the branch and the suspended bat dynamically interact through mechanical compliance [23].

To gain insight into the bat’s oscillation phenomena, the physical model (metronome) serves as an active analog of the bat system (Fig. 1 E, Movie 2, details in material and methods), while the 3D-printed bat model (Fig. 1 F) represents a passive mechanical counterpart. The phase-space analysis reveals a stable closed limit cycle (*ω ∼* 80 deg/s) for the active system, whereas the passive model exhibits damped spiral trajectories converging toward equilibrium (Fig. 1 H, I, fig. SI1F) and a maximum angular velocity *ω* of *∼* 100 deg/s reduces to zero.

The overlaid silhouettes of a bat (Fig. 1 J) reveals posture variability, highlighting dynamic body adjustments during roosting. The trajectory mapping (in Fig. 1K and Movie 1) demonstrates continuous oscillations over 50 s. The morphological descriptors (Fig. 1L) show average circularity, *ϕ* = 4 *π* A/P^2^ of 0.37 ± 0.03, while deformability, Γ = (a-b)/(a+b) remains lower at 0.30 ± 0.05, emphasizing adaptive reshaping strategies of bats, here A is area, P is perimeter, a is the major axis length, and b is minor axis length. The bats can adjust their grip from one to two to control oscillation during wind disturbance (Fig. S1J, Movie 3) and even stop oscillation. This reveals the bats are active oscillators, not just a passive hanging system. The acoustic analysis (Fig. 1M, Audio S1) reveals synchronized calls, characterized by dominant frequency peaks between 15–16 kHz.

## 2. Active Bats’ swaying under the wind

The field observations reveal a wind-driven transition from static hanging to synchronized swaying and escape behavior (Fig. 2A-B, fig. S2A-C, Movie 3). At moderate wind speeds, bats exhibit periodic angular oscillations with amplitude *θ ∼* 20–60^*°*^ and angular velocities *ω* up to *∼* 300 deg s^*−*1^, indicating forced pendulum-like dynamics (Fig. 2C-D, see in supplementary text section 2) and increased fluctuations preceding escape under strong wind forcing. As wind speed increases further, magnitudes in angular acceleration reach *α ∼* 10^3^–10^4^ deg s^*−*2^, (fig. S2D), destabilizing grip and triggering rapid runaway motion. This quantitative progression links aerodynamic forcing, collective synchronization, and biomechanical limits to bat detachment.

**FIG. 2.**
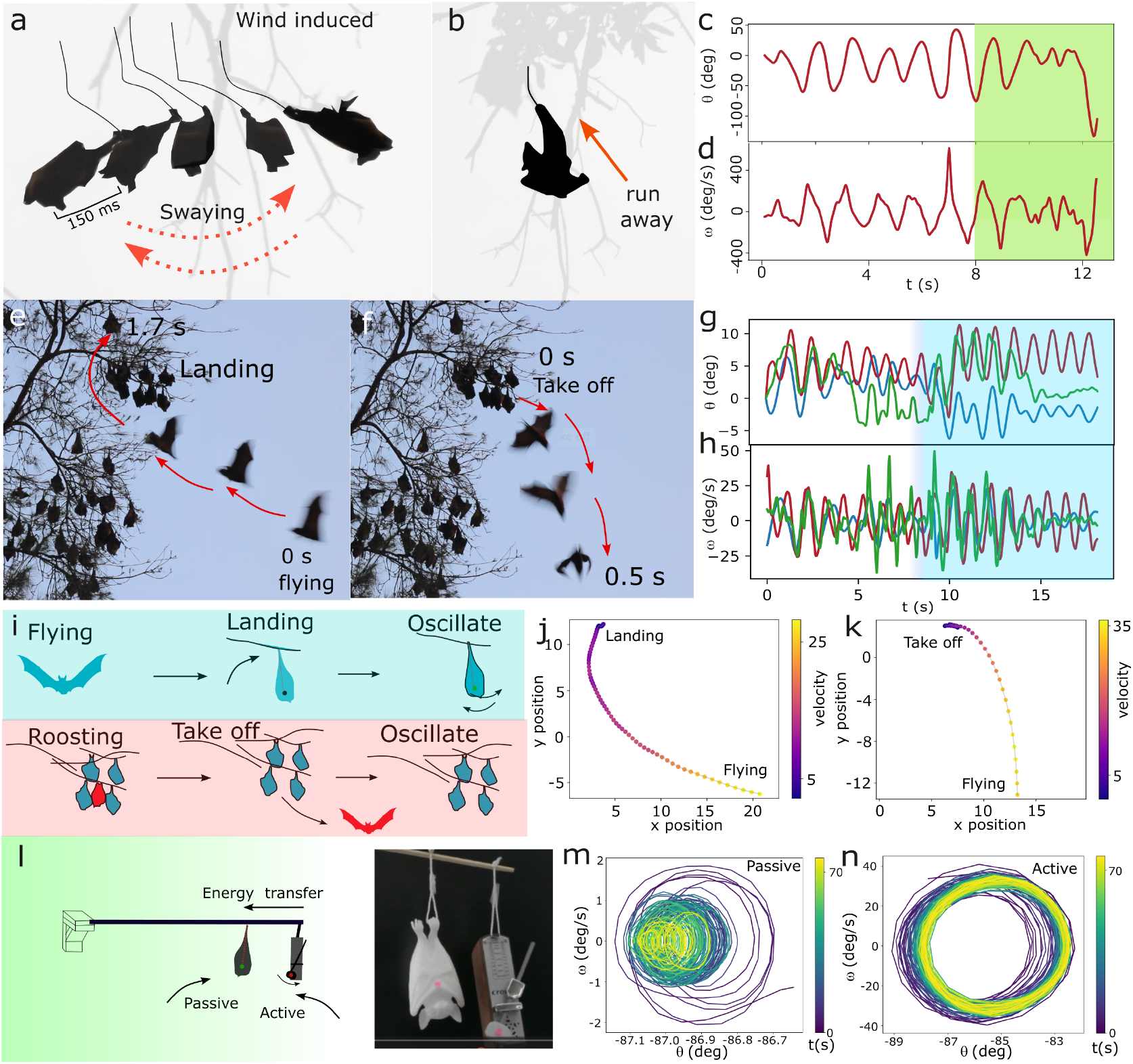
Bat oscillation under wind and externally induced force. (**a-b**) Bat swaying with high-speed wind with the moving branch. Afterward, it ran toward the branch’s joint. (**c**) Bat oscillating angle (*θ*) with progressing time. (**d**) Bat swaying angular velocity (*ω*). (**e**,**f**) Field image of the landing and take off of bats. (**g**,**h**) Bats take off, and landing induced the energy for oscillation represented in *θ* and *ω* with time for N=3. (**i**) Schematic of flying to landing, to induced oscillation, and take off to flying induced oscillation. (**j**,**k**) Bat landing and take-off trajectories with velocity overlay (bodydiameter/sec), where the diameter of the bats is used as a scale bar. (**l**) Schematic of hanging active and passive systems on a branch. (**m**,**n**) Phase-space (*ω* and *θ*) plots illustrating energy transfer from active to passive.

The individual bats approach the branch (Fig. 2E,J) from flight and gradually transition from active flapping to controlled landing to roosting on the tree. This process reflects an induced oscillation in the bat and branch. When a bat initiates flight (Fig. 2F,K), it pushes off the branch using its hind limbs and wings. This impulse transfers force to the flexible branch, causing a local deformation and oscillation of the branch structure and leading to small oscillatory motions in neighboring roosting bats (Fig. 2G,H).

The wind force follows aerodynamic drag, 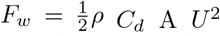, linking the observed deflection angle *θ* directly to wind velocity *U*, body shape, and exposed area, and enabling wind estimation from bats tilt measurements. The heatmaps demonstrate nonlinear growth of force with wind velocity and deflection angle, while increased mass reduces angular response under identical wind conditions (fig. S2H-J). When the wind is applied, the 3D-printed bat deforms by approximately 30 deg, being pushed by the airflow (fig. S2 E-G). Once the wind is turned off, the structure returns to its original position, indicating elastic recovery. However, in natural field conditions, wind forcing is often nonlinear and fluctuating, which can amplify the motion and induce stronger swinging dynamics in bats hanging from branches. The active system maintains a stable closed limit cycle, demonstrating continuous energy injection (Fig. 2L,N), while the passive system initially shows damped motion but gains energy from the active oscillator, expanding toward a synchronized stable limit cycle in phase space.

## 3. Two bats synchronization

When two bats hang upside down on a flexible shared branch, swaying collectively through mechanical coupling show in-phase and anti-phase synchronization (Fig. 3A, Movie 4). In-phase synchronization, the similar trajectories map show the synchronization behavior of bats (Fig. 3A,B, Movie 4). The speed of individual bats reaches a maximum of *∼* 15 body lengths per sec and closely overlaps, providing strong quantitative evidence of in-phase synchronization (Fig. 3C). For anti-phase synchronization, the trajectory maps shown highlight movements over time (20 s), color-coded by seconds (Fig. 3D,E), revealing synchronized oscillations and dynamic coordination that emerge naturally during roosting behavior. The phase difference reflects the anti-phase synchronization after 10 s (Fig. 3H). The phase-space trajectories reveal that both the branches and bats undergo coupled oscillations (angular speed *ω ∼* 40 deg/s and angular acceleration *α ∼* 400 deg/s), with the bats displaying large amplitudes and synchronization under midden wind influence (Fig. 3F,G,I and fig. S3A-C). For two bats hanging on the shared flexible branch, the elastic restoring torque 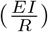 balances the gravitational torque 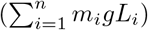, forming a mechanically coupled series elastic system [24].

**FIG. 3.**
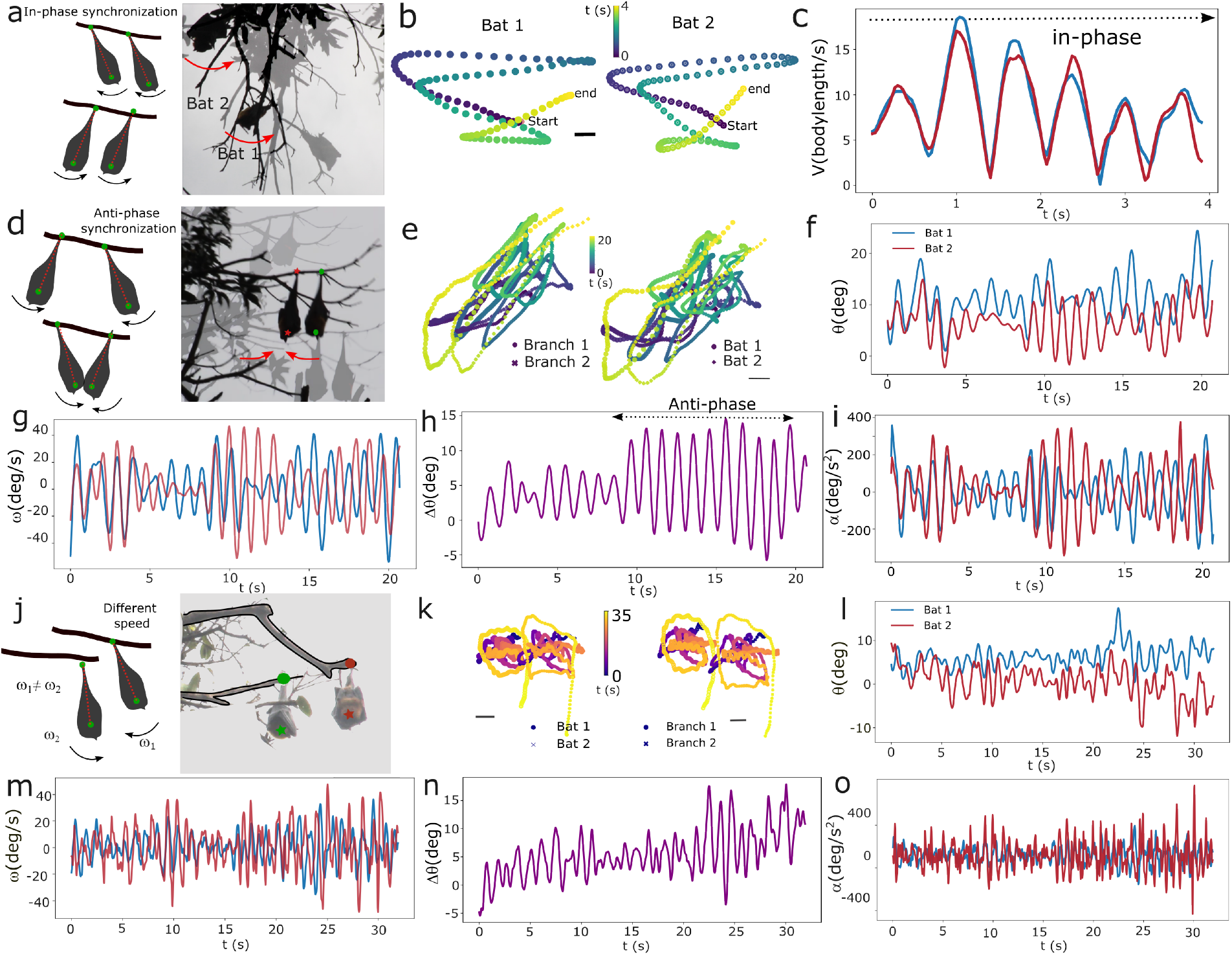
Two bats oscillations on the shared and separated branches. (**a**) Schematics and field image of the inphase synchronous state of two bats. (**b**,**c**) In phase synchronization of two bats’ trajectories and speed, V (bodylength/s). (**d**) Schematics and field observation of the anti-phase synchronous state of bats roosting on tree branches, overlaid with the traced branch and bat positions (N=2). (**e**) Time-resolved trajectories of bats and branch tracked over 20 s, where the color bar represents progression from 0 (purple) to 20 s (yellow). (**f-i**) Angular displacement *θ*, angular velocity *ω*, phase shift difference Δ*θ*, angular acceleration 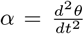 plot with time. (**j**) Schematic trajectories of bats swaying on separated flexible branches and field observation of bats roosting on tree branches, overlaid with the traced branch and bat positions. (**k**) Time-resolved trajectories of bats and branch tracked over 35 s, where the color bar represents progression from 0 (purple) to 35 s (yellow). (**l-o**) Angular displacement *θ*, angular velocity *ω*, phase shift difference Δ*θ*, and angular acceleration *α* plot with time.

The swaying dynamics of bats hanging from separated thin, flexible branches reveal intricate motion patterns driven by environmental forces, such as wind (Fig. 3J, Movie 4). These branches allow bats to oscillate in response to external perturbations, as shown in the trajectories of Fig. 3K. The angular velocity *ω* and acceleration *α* of individual bats are different magnitudes, and the phase difference (Fig. 3L-O) displays nonlinear dynamics, leading to non-synchronized collective behavior. For bats swaying on a separated branch, a simple torque balance between elastic 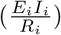 and gravitational moments gives (m_*i*_ g L_*i*_) for each branch.

## 4. Active oscillators synchronization

We performed the experiment with two active oscillators to replicate the two bat swaying observations (Movie 5). The two active oscillators show the beat synchronization with two phases, which are in-phase and anti-phase synchronization (Fig. 4A,B). The synchronization of pair active oscillators on a shared branch is highly dependent on the material’s rigidity and its deformation capacity. When oscillators are placed on a stiff metal branch, the lack of physical deformation results in weaker coupling, leading to a slower synchronization process that yields only 4 beat cycles in 70 seconds(Fig. 4G left panel). Conversely, a soft wooden branch deforms during oscillation; this mechanical flexibility facilitates a faster transition to synchrony, achieving 5 beat cycles in the 70-second time frame (green line). The two oscillators do fast beat synchronizations when they are close together (6 cm), but increasing the distance by around 3 times between them, slows down the synchronization process and doubles (T *∼* 30s) the beat period (Fig. 4G right panel). The active oscillators with different frequencies do not show any phase locking or synchronization. When oscillators are on separate branches, Δ*θ* = 40–60^*°*^ and no limit-cycle overlap, indicating negligible coupling (Fig. 4C,D). The beating period (*T*) can be defined by the function *T* (*ρ, E, d, v*) where *d* is the distance, *E* is the Young’s modulus of the branch, *ρ* is the branch density, and *v* is the speed of the branch.

**FIG. 4.**
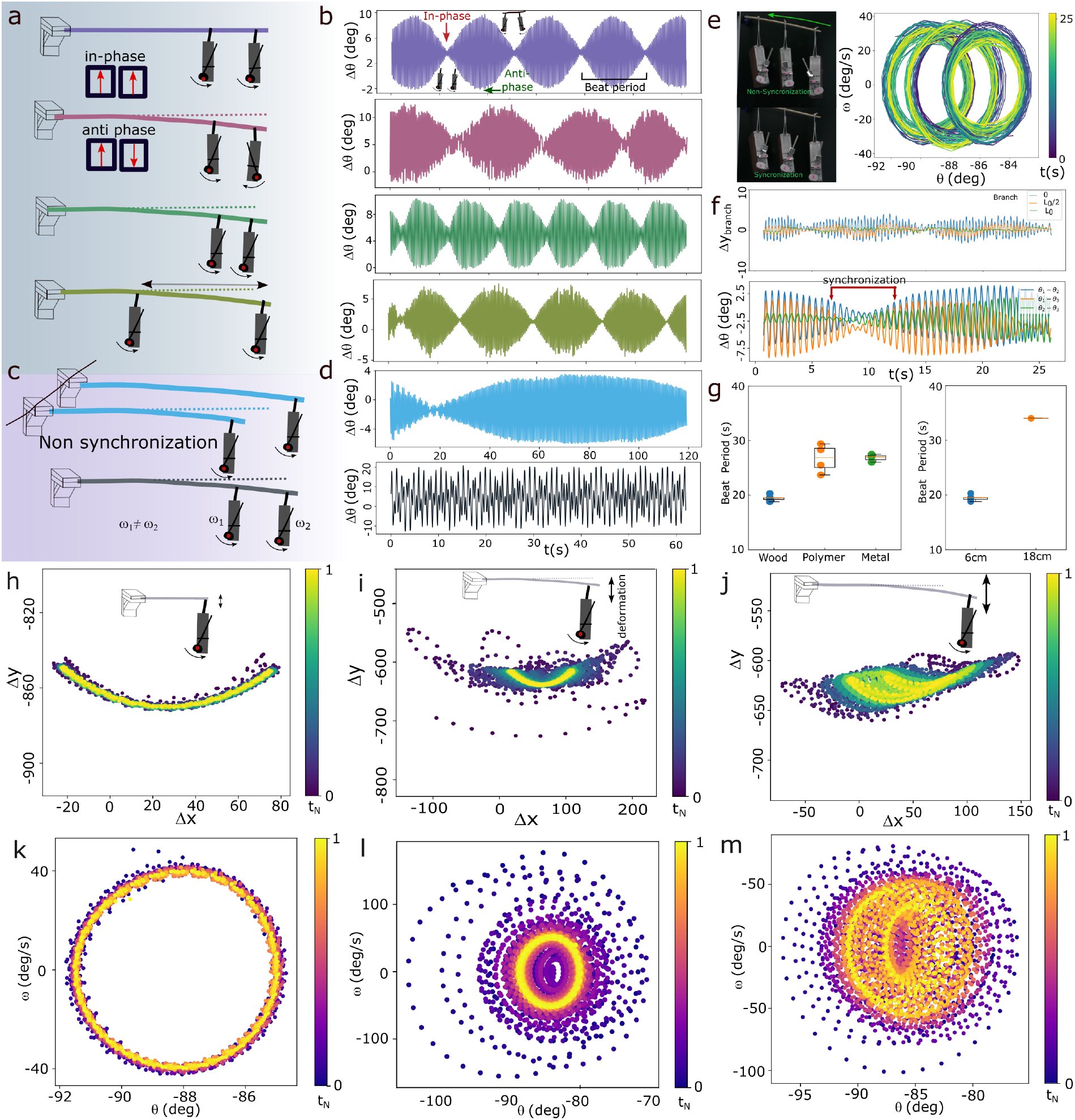
Two physical active oscillators synchronization. (**a**,**b**) Schematic of a two-oscillating active system. The phase differences (Δ*θ*) among active system elements (metal, polymer, and wood) and different distances for the wood rod. (**c**,**d**) Schematic of a non-synchronization oscillating active system and Δ*θ* for the separated branch (sky blue line) and different frequencies (black line) of oscillators. (**e**) Schematic of three active systems and the phase-space (*θ, ω*) of all three oscillators. Three active oscillators Δ*θ* with time. (**f**) The branch vertical deformation (Δy), phase difference (Δ*θ*_12_,Δ*θ*_13_,Δ*θ*_23_) between the active oscillators. (**g**) Box plots represent the beat period (T, sec) influenced by branch materials (for 6 cm) and distance (for 6 cm to 18 cm) between oscillators for wood. (**h-j**) Induced the branch vertical deformation (Δy), the trajectories of active oscillators where the branch is the reference point for shorter to longer branches. (**k-m**) Phase space between *ω* and *θ* for shorter to longer branch.

To understand the multiple active oscillator behavior in comparison to two on the same branch, the three active oscillators reduce the phase difference from *∼* 50^*°*^ to *<* 5^*°*^, and the frequency mismatch approaches zero, demonstrating strong mechanical coupling through the shared compliant branch (Fig. 4E-G). The shift from extending the branch length enhances structural compliance, leading to more deformations (Movie 5). This physical change shifts the spatial trajectories from constrained, narrow arcs into broader (Fig. 4H-M, fig. S4A-C). Similarly, the phase-space topography transitions from circular limit cycles to increasingly dispersed structures, signaling a rise in nonlinear coupling and a more sophisticated level of oscillatory complexity within the oscillating branch.

## 5. Collective active bats’ Oscillation

The multiple bats are distributed around a branching tree during roosting, captured phase-colored angular dynamics (*θ*) and synchronization patterns relative to a shared oscillating branch (Fig. 5A, Fig. S5A-B, Movie 6), where individual oscillators (*N* = 11) exhibit natural angular velocities (*ω*_0_). Bats and moving branch (gray line) trajectories are shown in (*x, y*) coordinates with overlay time, revealing stable limit cycles (Fig. 5B). The pairwise mean phase difference matrix, Δ*θ*, of bats in a heat map (Fig. 5C), with small phase lags dominating where structured patterns suggest cluster coordination without a dominant leader, indicating distributed mechanical coupling. The mean angular position (*θ*, red curve, Fig. 5D) reveals periodic oscillations corresponding to a peak-to-peak amplitude for eleven bats, indicating stable limit-cycle dynamics. The phase difference density map (Fig. 5E) demonstrates the collective synchronization among the 11 coupled oscillators, with phase differences Δ*θ* clustering near 0°, indicating stable frequency and phase locking over a 15 s duration. The collective exhibits limit-cycle oscillations with a mean angular velocity of *ω ∼* 18 deg/s, the small periodic velocity fluctuations (Fig. 5F) confirm a symmetric forward-backward motion with bounded dynamics. The difference in periods (Δ*T* = *T*_*i*_ *− T*_*j*_) is prominent (Fig. 5 G) when bats are distantly related (*i* = 11 and *j* = 3). In contrast, when bats occupy the same branch (*i* = 11 and *j* = 7), they exhibit very similar periods (bats 11 and 7). The system exhibits characteristic features of weakly coupled non-linear oscillators in which the coupling-to-damping ratio determines the synchronization threshold. We also analyzed the oscillation of a large group of bats (details in the supplementary text, Fig. S5C-G), where the spatial trajectories of the colony, along with the velocity components (*v*_*x*_, *v*_*y*_), revealed that strong wind conditions generate complex, chaotic, and collective oscillatory dynamics within the roosting group.

**FIG. 5.**
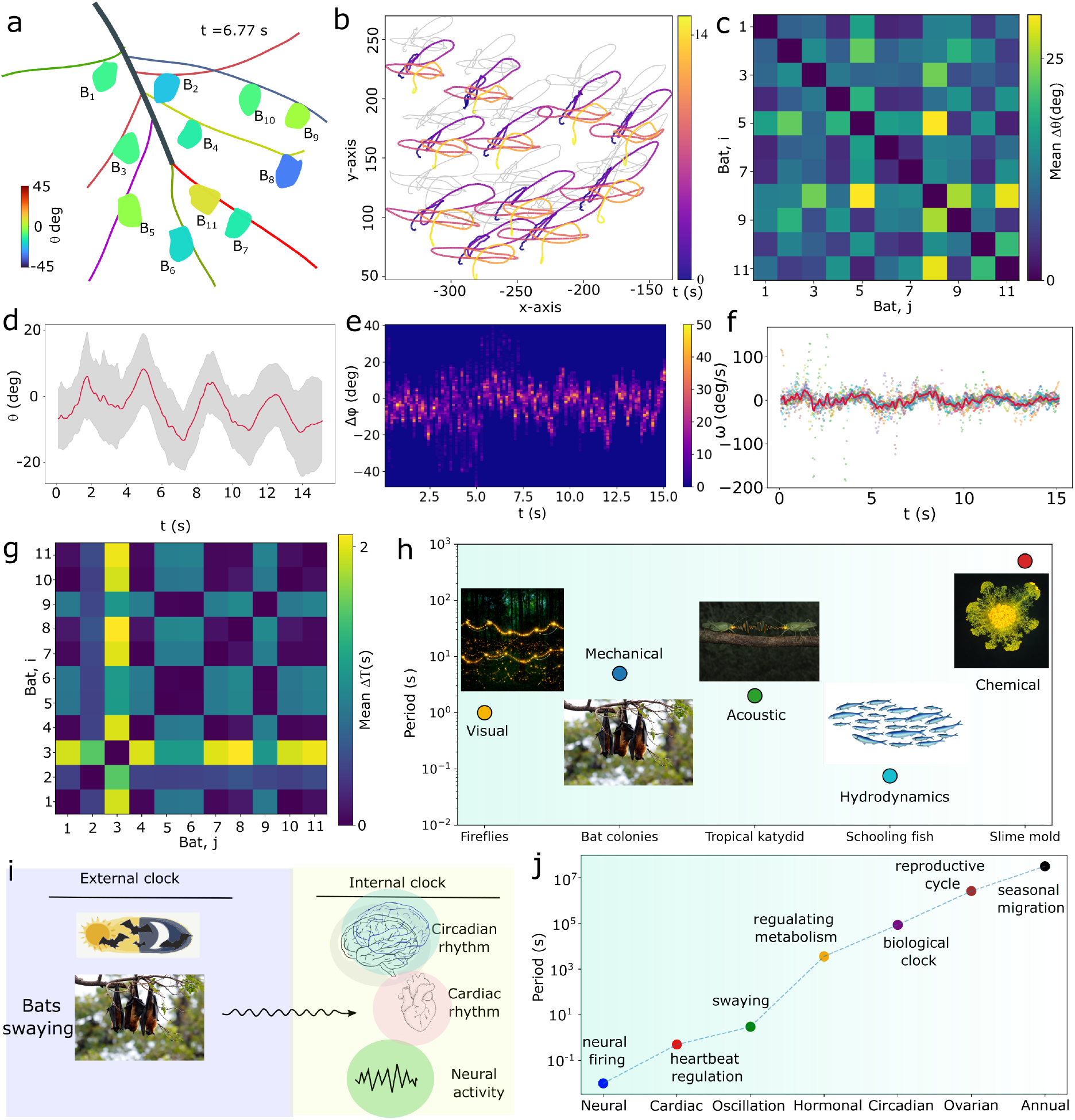
Collective active bats oscillation. (**a**) Field snapshot of a tree canopy with multiple hanging markers highlighting individual bats roosting on branches. Each marker corresponds to a tracked individual used for motion analysis of the group synchronization of hanging bats. (**b**) Plan-view trajectories (x,y) of individual bats, color-coded by identity, showing localized oscillatory motion with limited net displacement across the canopy. (**c**) Bats oscillator (i,j) phase difference (Δ*θ*). (**d**) The mean angular position (*θ*) of the oscillatory system as a function of time (black line), with the shaded gray region representing standard deviation across oscillators. (**e**) Spatiotemporal density map phase difference (Δ*θ*) with time (t). (**f**) The colored scatter points show individual bats’ oscillator velocities, while the red curve indicates the mean angular velocity (*ω*) for N=11 bats. (**g**) Bats oscillator (i,j) period difference (ΔT). (**h**) Comparison between different animal synchronization systems. (**i**,**j**) Biological rhythm of bats across temporal scale.

## DISCUSSION

Synchronization occurs in nature with various sensory modes, such as fireflies with visual, tropical katydids with acoustic, slime molds with chemical, and bat colonies with mechanical coupling across nature (Fig. 5H, Table S1) [25]. The external and internal biological clocks of bats span ten orders of magnitude (10^*−*1^ to 10^7^ s), and bats’ oscillation occurs with *∼* 1 s, same as the neural timescale regime (Fig. 5I,J, Table S2). The collective swaying of bats roosting on flexible tree branches represents a macroscopic oscillation dynamic through mechanical rather than biochemical or synaptic interactions.

Bats behave as active self-oscillators and actively alter body posture (stretching wings, shifting grips, curling), which changes the aerodynamic and inertial profile [26]. Bats are vocalizing during swaying, and their sounds could serve as communication signals, either distress calls or coordination calls [27], which are toward active behavioral responses. The bat’s hippocampus plays a broader role than simply mapping an individual’s location; its neurons capture details about neighboring bats, from their identities to their proximity [28, 29]. Some bat species exploit wind conditions and harness advancing storm fronts to maintain long-distance migrations [30]. During roosting, the wind waves pass through the tree canopy [31–36] and energy is transferred to induce the bat’s oscillation.

The physical active oscillators, compared with biological bats colonies, exhibited idealized oscillatory behavior due to their controlled mechanical properties and less biological variability. The dynamics of physically coupled active oscillators observed beat oscillations emerging from their mechanical coupling. The beat patterns were influenced by the flexibility of the supporting material and the distance-dependent coupling between oscillators, with compliant substrates enhancing energy transfer and synchronization. Bats’ display complex swaying dynamics arising from modulating grip control, body reorientation, heterogeneous natural frequencies, and continuous responses to environmental perturbations. Many animals adopt the biomechanical behavior of trees [37]. Here, we characterize flying fox bats as active oscillatory systems in which individuals in both small and large groups exhibit collective swaying behavior that may synchronize via mechanical coupling. In the future, we can conduct comparative studies between bat species on how morphology, habitat selection, and colony density have shaped the evolution of synchronized swaying motion and the transition between order and chaos.

## References

[1] J. T. Emlen, Flocking behavior in birds, The Auk 69, 160 (1952).

[2] J. Buck and E. Buck, Biology of synchronous flashing of fireflies, Nature 211, 562 (1966).

[3] D. H. Cushing and F. R. Harden Jones, Why do fish school?, Nature 218, 918 (1968).

[4] E. T. Nelson, Swarming insects, Science ns-4, 111 (1884).

[5] O. Peleg, J. M. Peters, M. K. Salcedo, and L. Mahadevan, Collective mechanical adaptation of honeybee swarms, Nature Physics 14, 1193 (2018).

[6] I. R. Epstein, K. Kustin, P. De Kepper, and M. Orbán, Oscillating chemical reactions, Scientific American 248, 112 (1983).

[7] I. D. Couzin, Synchronization: the key to effective communication in animal collectives, Trends in cognitive sciences 22, 844 (2018).

[8] P. K. Bhattarai, B. Sharma, B. Bhattarai, and B. Neupane, Spatio-temporal colony dynamics and roost tree preferences of indian flying fox, pteropus medius, in jhapa, nepal, Biotropica 57, e70130 (2025).

[9] D. B. Boerma and S. M. Swartz, Roosting ecology drives the evolution of diverse bat landing maneuvers, iScience 27, 110381 (2024).

[10] N. Sungar, J. Sharpe, L. Ijzerman, and J.-W. Barotta, Synchronization and self-assembly of free capillary spinners, Physical Review E 111, 035104 (2025).

[11] J.-W. Barotta, G. Pucci, E. Silver, A. Hooshanginejad, and D. M. Harris, Synchronization of wave-propelled capillary spinners, Physical Review E 111, 035105 (2025).

[12] H. M. Oliveira and L. V. Melo, Huygens synchronization of two clocks, Scientific reports 5, 11548 (2015).

[13] C. Huygens, Horologium oscillatorium (1980).

[14] G. Neuweiler, The biology of bats (Oxford University Press, 2000).

[15] D. Howell and J. Pylka, Why bats hang upside down: a biomechanical hypothesis, Journal of Theoretical Biology 69, 625 (1977).

[16] T. H. Quinn and J. J. Baumel, Chiropteran tendon locking mechanism, Journal of Morphology 216, 197 (1993).

[17] W. Schutt, Jr, Digital morphology in the chiroptera: the passive digital lock, Cells Tissues Organs 148, 219 (1993).

[18] S. Mallick, A. Hossain, and S. K. Raut, On the significance of bats’ hanging posture, International Journal of Research and Analytical Reviews (IJRAR) 9 (2022), available at: www.ijrar.org.

[19] V. Elangovan, Effect of tree characteristics on roost selection of the indian flying fox, pteropus giganteus, Journal of Bat Research & Conservation (2019).

[20] B. Murugavel, S. Kandula, H. Somanathan, and A. Kelber, Home ranges, directionality and the influence of moon phases on the movement ecology of indian flying fox males in southern india, Biology Open 12, bio059513 (2023).

[21] J. W. Kanwisher, G. Gabrielsen, and N. Kanwisher, Free and forced diving in birds, Science 211, 717 (1981).

[22] D. G. Raveling, Migration reversal: a regular phenomenon of canada geese, Science 193, 153 (1976).

[23] B. A. Gardiner, A dynamic model of the behaviour of sitka spruce in high winds, Tree Physiology 10, 101 (1992).

[24] E. Virot, A. Ponomarenko, É. Dehandschoewercker, D. Quéré, and C. Clanet, Critical wind speed at which trees break, Physical Review E 93, 023001 (2016).

[25] A. Goldbeter, Biological rhythms: clocks for all times, Current biology 18, R751 (2008).

[26] D. B. Boerma et al., Wings as inertial appendages: How bats recover from disturbances, Journal of Experimental Biology 222, jeb204255 (2019).

[27] M. C. Rose, B. Styr, T. A. Schmid, J. E. Elie, and M. M. Yartsev, Cortical representation of group social communication in bats, Science 374, eaba9584 (2021).

[28] E. D. Roth, A. J. Yu, S. J. Y. Mizumori, et al., Natural switches in behaviour rapidly modulate hippocampal coding, Nature 608, 555 (2022).

[29] A. Forli and M. M. Yartsev, Hippocampal representation during collective spatial behaviour in bats, Nature 621, 796 (2023).

[30] E. Hurme, I. Lenzi, M. Wikelski, T. A. Wild, and D. K. N. Dechmann, Bats surf storm fronts during spring migration, Science 387, 97 (2025).

[31] J. Moore, B. Gardiner, and D. Sellier, Tree mechanics and wind loading, in Plant biomechanics: from structure to function at multiple scales (Springer, 2018) pp. 79–106.

[32] K. James, Dynamic loading of trees, Arboriculture & Urban Forestry (AUF) 29, 165 (2003).

[33] K. R. James, N. Haritos, and P. K. Ades, Mechanical stability of trees under dynamic loads, American journal of Botany 93, 1522 (2006).

[34] T. Jackson, A. Shenkin, A. Wellpott, K. Calders, N. Origo, M. Disney, A. Burt, P. Raumonen, B. Gardiner, M. Herold, et al., Finite element analysis of trees in the wind based on terrestrial laser scanning data, Agricultural and Forest Meteorology 265, 137 (2019).

[35] H.-C. Spatz and B. Theckes, Oscillation damping in trees, Plant science 207, 66 (2013).

[36] L. Tadrist, M. Saudreau, P. Hémon, X. Amandolese, A. Marquier, T. Leclercq, and E. De Langre, Foliage motion under wind, from leaf flutter to branch buffeting, Journal of the Royal Society Interface 15, 20180010 (2018).

[37] N. H. Hunt, J. Jinn, L. F. Jacobs, and R. J. Full, Acrobatic squirrels learn to leap and land on tree branches without falling, Science 373, 697 (2021).

